# A novel mouse model of focal limbic seizures that reproduces behavioral impairment associated with cortical slow wave activity

**DOI:** 10.1101/2021.05.05.442811

**Authors:** Lim-Anna Sieu, Shobhit Singla, Abdelrahman Sharafeldin, Ganesh Chandrasekaran, Marcus Valcarce-Aspegren, Ava Niknahad, Ivory Fu, Natnael Doilicho, Abhijeet Gummadavelli, Cian McCafferty, Richard Crouse, Quentin Perrenoud, Marina Picciotto, Jessica Cardin, Hal Blumenfeld

## Abstract

Patients with focal temporal lobe seizures often experience loss of consciousness. In humans, this loss of consciousness has been shown to be positively correlated with EEG neocortical slow waves, similar to those seen in non-REM sleep. Previous work in rat models of temporal lobe seizures suggests that decreased activity of subcortical arousal systems cause depressed cortical function during seizures. However, these studies were performed under light anesthesia, making it impossible to correlate behavior, and therefore consciousness, to electrophysiologic data. Further, the genetic and molecular toolkits allowing for precise study of the underlying neural circuitry are much more developed in mice than in rats. Here, we describe an awake-behaving, head-fixed mouse model of temporal lobe seizures with both spared and impaired behavior reflecting level of consciousness. Water-restricted mice were head-fixed on a running wheel and trained to associate an auditory stimulus to the delivery of a drop of water from a dispenser. To investigate the effect of seizures on behavior, seizures were electrically induced by stimulating either the left or right hippocampus via a chronically-implanted electrode, while mice were performing the task. Behavior was measured by monitoring lick responses to the auditory stimulus and running speed on the wheel. Further, local field potentials (LFP) signals were simultaneously recorded from hippocampus and orbitofrontal cortex (OFC). Induced focal seizures were 5-30s in duration, and repeatable for several weeks (n=20 animals). Behavioral responses showed a decrease in lick rate to auditory stimulus, and decreased running speed during seizures (p<0.01, n=20 animals). Interestingly, licking response to sound could vary from being impaired to normal during seizures. We found that behavioral impairment is correlated with large amplitude cortical slow-wave activity in frontal cortex, as seen in patients with temporal lobe seizures. These results suggest that induced focal limbic seizures in the mouse can impair consciousness and that the impaired consciousness is correlated with depressed cortical function resembling slow wave sleep. This novel mouse model has similar characteristics with previously studied rat models and human temporal lobe seizures. By leveraging the genetic and molecular techniques available in the mouse, this model can be used to further uncover fundamental mechanisms for loss of consciousness in focal seizures.

## Introduction

Temporal lobe epilepsy (TLE) is one of the most common and debilitating forms of epilepsy in patients with intractable seizures (de Boer et al. 2008; Charidimou and Selai 2011). They are characterized by focal seizures originating from limbic structures including the hippocampus. These seizures often cause functional deficits far beyond those expected from local temporal lobe impairment, including loss of consciousness. Interestingly, patients with temporal lobe seizures with impaired consciousness show slow oscillations (1-2 Hz) in frontal and parietal cortices, similar to a state resembling deep sleep, anesthesia, and coma. The seizures themselves are confined to the temporal lobe, particularly in limbic structures (Blumenfeld et al. 2004b). Human studies suggest that focal temporal lobe seizures depress cortical function that lead to impaired consciousness not through direct seizure propagation, but rather by reducing subcortical arousal, causing slow-wave sleep-like state in the cortex (Blumenfeld et al. 2004a; Englot et al. 2010). The mechanism by which the seizures, confined to a brain region devoted to memory, emotions, and related functions, can cause depressed cortical function leading to loss of consciousness have yet to be fully understood. Rat models recapitulate key components of ictal unconsciousness in humans, including transition from waking rhythms into slow-wave activity in the frontal cortex during focal limbic seizures (Englot et al. 2008; Englot and Blumenfeld 2009; Motelow et al. 2015). Studies in rat cortical slow wave activity observed during those seizures showed alternating up states of increase firing and down states of quiescence (Englot et al. 2008; Yue et al. 2020), similar to natural sleep or deep anesthetic conditions (Steriade et al. 1993b; Steriade et al. 2001; Haider et al. 2006). Furthermore, cortical and thalamic cholinergic neurotransmission, critical for maintaining arousal state (Steriade et al. 1993a; Hill and Tononi 2005; Munoz and Rudy 2014), is reduced during seizures (Motelow et al. 2015), notably attributable to a decrease of cholinergic neuronal firing activity from basal forebrain and the pedunculopontine tegmental nucleus (PPT) (Motelow et al. 2015; Andrews et al. 2019). Moreover, other key regions in subcortical arousal systems, such as central lateral nucleus of thalamus and the paratenial nucleus of the thalamus (Van der Werf et al. 2002; Schiff et al. 2013) showed reduced neuronal firing activity (Feng et al. 2017; Zhao et al. 2020). Meanwhile, GABAergic systems known to be strongly connected to the hippocampus, such as lateral septum (LS) or anterior hypothalamus (Risold and Swanson 1997; Cenquizca and Swanson 2006), showed increase blood oxygen level dependent (BOLD) (Motelow et al. 2015), suggesting an activation of inhibitory structures that depress subcortical arousal systems during focal limbic seizure. Further studies revealed that electrical stimulation of LS produced a transition to cortical slow wave activity with reduced cortical choline neurotransmission in the absence of seizures (Li et al. 2015). These findings strongly support the hypothesis that subcortical arousal systems are suppressed during focal limbic seizures, causing reduced arousal output to the cortex and impaired consciousness, possibly from activation of GABAergic systems such as the LS inhibitory inputs.

However, these studies on the rat model were performed under light anesthesia, which can alter the cortical and subcortical physiology, as well as the network effects of stimulation (Alkire et al. 2008). Further, due to the anesthetize state, results could not be directly linked to behavior. Therefore, we sought to investigate network, neurotransmitter and neuronal mechanisms responsible for impaired consciousness during focal limbic seizures in a new awake and behaving animal model. Due to a broader availability of genetic and molecular tools and the ease to manipulate mice in an awake head-fixed condition, we developed an awake, behaving head-fixed mouse model of TLE to measure electrophysiological and behavioral effects of focal limbic seizure. This mouse model will provide new opportunities to identify neuronal mechanisms crucial for maintaining consciousness during focal limbic seizures and explore the possibility to restore cortical function and behavior (Gummadavelli et al. 2015; Furman et al. 2015; Kundishora et al. 2017; Xu et al. 2020), providing hope for new treatment approaches to restore arousal in human epilepsy. This novel animal model of focal limbic seizures has been described recently by our group in abstract form and is presented here in greater detail (Sieu et al. 2020; Sieu et al. 2018; Sieu et al. 2019).

## Methods

### Animal Preparation and Surgery

All procedures were performed in accordance with approved protocols of Yale University’s Institutional Animal Care and Use Committee. Adult female C57BL/6 mice at 3-6 months of age were used in these experiments. All surgeries were performed under deep anesthesia with Ketamine (90 mg/kg) and Xylazine (9 mg/kg). During surgery, a titanium head-plate was attached to the animal’s skull with dental cement and bipolar electrodes were placed in the orbitofrontal cortex (OFC) and bilateral hippocampus (HC). Mice were allowed a week of recovery before starting auditory task training session followed by behavioral experiments. All recordings occurred in awake, head-fixed animals running freely on a wheel. Once experiments were completed, animals were euthanized. Brains were then harvested for histological analysis.

### Auditory task

Mice were allowed to recover for one week and then water restricted in order to promote appetite response to liquid reward during behavioral study and awake recordings. Animals were then acclimated to the experimental set up including head-fixation and running wheel. They were then trained to lick from a lick port containing a drop of water (5μL) in response to an auditory stimulus. The auditory stimulus occurred every 10-15s during the training session. One training session lasted for a minimum 10min or until the animal stopped licking successive drops. Behavioral experiments started after the animal reached 95-100% success of licking water after the stimulus, which happened approximately after 2-3 days of training.

During behavioral experiments, the auditory stimulus followed by water presentation occurred every 10-15s during a baseline period of at least 60s before a seizure was induced. During seizures, the auditory stimulus was emitted every 3-7s interval. During post-ictal and recovery period, the stimuli were back to 10-15s interval. The behavioral experiment session ended when the animal stopped licking successive drops or when the session reached a maximum of 2 hours. Mice underwent behavioral experiments 5 days a week for a duration of maximum 2 months. During those periods, mice were weighed daily and water restricted to reach up to18% of weight loss, with free access to food.

### Electrophysiology

#### Seizure induction and cortical recordings

Seizure initiation and local field potential (LFP) recordings were performed via bilaterally chronic-implanted bipolar electrodes in the dorsal hippocampi. Additional cortical LFP recordings were taken via a chronic-implanted bipolar electrode in the right orbitofrontal cortex (OFC). Focal seizures were induced in one side of the hippocampus (HC) using a 2s, 60 Hz current injection. One session of recording included both electrophysiology and behavioral experiments (licking and/or running). Seizures were triggered once per session and per day for a maximum of two months. Sham control stimulations without seizures in HC were also conducted before seizure stimulation or after seizure recovery. The LFP signals collected by the bipolar electrodes was filtered at 0.1-100 Hz and 1-500 Hz, for OFC and HC respectively, and were acquired as described previously (Englot et al. 2008; Englot and Blumenfeld 2009; Schridde et al. 2008). Data were analyzed using Spike2 and MATLAB. Baseline was defined as a period of 60s before seizure, post-ictal as the 30s period after seizure, and recovery as a 60s period after post-ictal.

## Results

### Unilateral electrostimulation in dorsal hippocampus induces focal limbic seizures with reduced behavioral response

Performing a behavioral task requires a minimum of arousal and engagement. Therefore, behavioral responsiveness measured here was used as an indicator of level of consciousness. To determine if behavioral responses were affected by focal limbic seizures in mice, seizures were induced by electrically stimulating unilateral dorsal HC (Englot et al. 2008), while they performed an auditory task on a head-fixed running wheel apparatus (Fig 1A). Induced seizures lasted between 5-30 s, with an average of 15.3±6.5 s, and were repeatable for several weeks (n=20 animals). The average duration of a session was 18.3±8.6 min before mice became satiated to the water reward and stopped responding. HCs and OFC LFP signals, sound presentations, lick responses and wheel position were simultaneously recorded (Fig1.B).

**Figure 1.**
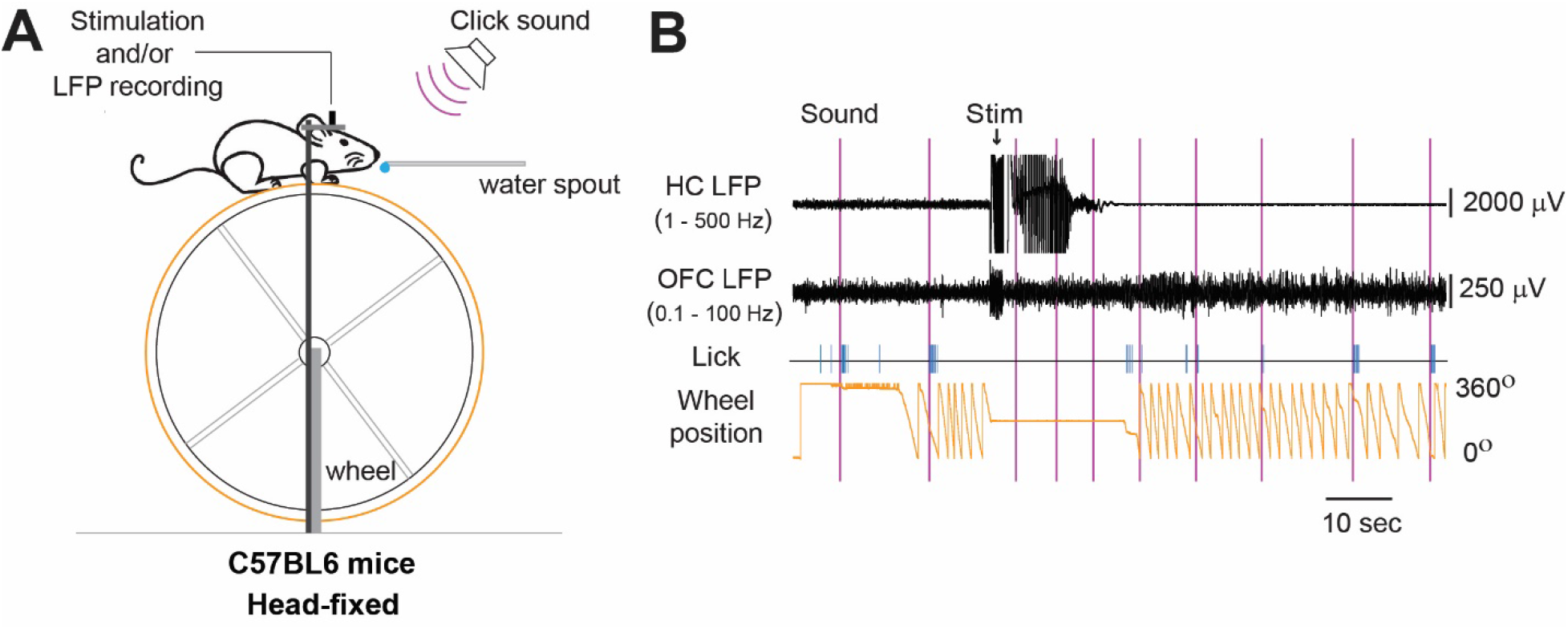
Experimental apparatus and recording set up. A. Mice were head fixed and allowed to run on a running wheel while simultaneously obtaining recordings from the hippocampus (HC) and orbitofrontal cortex (OFC). An auditory stimulus indicated water was available through a lick spout. B. Example trace from a single experiment. Auditory stimuli are represented by vertical pink lines. Arrow signifies stimulus artifact from electrical stimulus causing seizures. Note that the animal stopped running (indicated by no change in wheel position in bottom trace), and stopped licking to the auditory stimulus (as indicated by the vertical blue traces) during the ictal period.

During the baseline period, mice responded to the auditory stimulus by licking 99.5% (854/858 stimuli) of the time. The first lick response occurred on average 137±161.0 ms after the sound, with a mean lick rate of 10.5±1.6 licks per second. Induction of a focal limbic seizure showed significantly reduced responsiveness to auditory stimuli during ictal period compared to baseline, and during post-ictal and recovery period but with a tendency to return back to normal (Fig2.A&B). Sham stimulation without seizure did not evoke any significant difference in behavioral responses compared to baseline period (data not show), suggesting that impaired behavior observed was caused by seizures, not stimulation itself. Interestingly, lick response to sound could vary from being impaired to normal during the ictal period (Fig2.A, B&C), suggesting that consciousness is not always impaired during seizures, an observation also seen in humans. Moreover, analysis of wheel speed showed that mice are significantly more immobile during the ictal period, and then ran significantly more during the recovery period compared to baseline (Fig2.D), supporting the idea that seizures can affect overall behavior in mice. These results suggest that induced focal limbic seizures in mice are associated with both impaired and spared behavior, as seen in patients with TLE.

**Figure 2.**
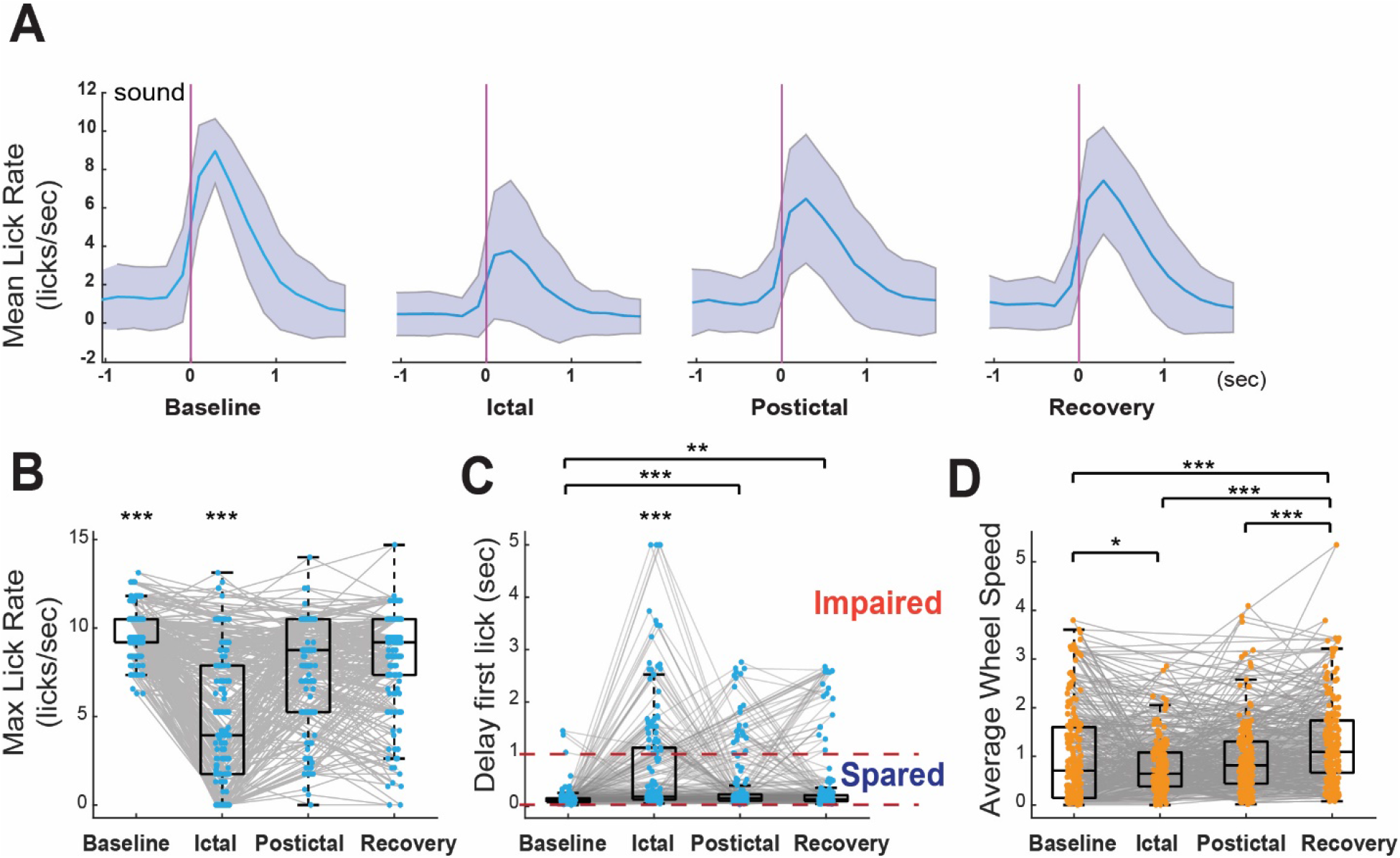
Induced focal limbic seizures in mice cause impaired behavior. A. Mean lick rates (blue trace) following sound presentation (vertical pink trace) are compared at different periods: baseline, ictal, postictal and recovery. Ictal period showed a decreased lick rate compared to baseline. B&C. Scatter boxplot showing average of maximum lick rate (B) and average of delay to the first lick (C) following sound presentation during each period for each recording. A grey line linking the dots at different periods represents one recording. Seizures significantly decrease lick rate (B) and significantly increase latency to first lick (C). Horizontal dashed line in C indicates lick latency of 1s, with lick latencies above 1s categorized as impaired behavior. D. Scatter boxplot showing average wheel speed for each period for each recording. Mice showed significantly decreased movement during seizures. Error bars show +/-standard deviation. ANOVA statistic test was used for multigroup comparison, n=20 animals, * p<0.05; ** p<0.02; *** p<0.01.

### Cortical slow waves during induced partial limbic seizure is associated with impaired behavior

In patients, temporal lobe seizures with impaired consciousness show a transition from waking rhythms into slow wave oscillations (1-2Hz) in frontal cortex, resembling sleep. In contrast, temporal lobe seizures without loss of consciousness are not accompanied by this change (Blumenfeld et al. 2004a; Englot et al. 2010). We sought to determine if our mouse model of TLE reproduced this characteristic. In mice, the frequency of spontaneous slow wave sleep oscillates between 1-4Hz. To correlate OFC LFP activity to behavior, we classified behavioral responses into two categories: impaired behavior corresponding to impaired consciousness and spared behavior for unimpaired consciousness based on first lick latency to the auditory stimulus: ‘s*pared*’ = first lick occurred within 1s window after the auditory stimulus, ‘*impaired*’ = no lick or first lick occurred after 1s window after the auditory stimulus. Power spectrum was calculated for OFC LFP signals within the time windows of 2s before and 2s after each classified sound. We found that induced focal limbic seizures could evoke slow wave activity in the OFC (Fig.3A), and that, cortical slow wave activity was strongly associated with impaired behavior in contrast with spared behavior (Fig. 3B&C), which replicates human findings. These results demonstrated that our new mouse model shares characteristics with the rat model and patients with temporal lobe epilepsy and adds the possibility to assess behavior related to neuronal network physiology.

**Figure 3.**
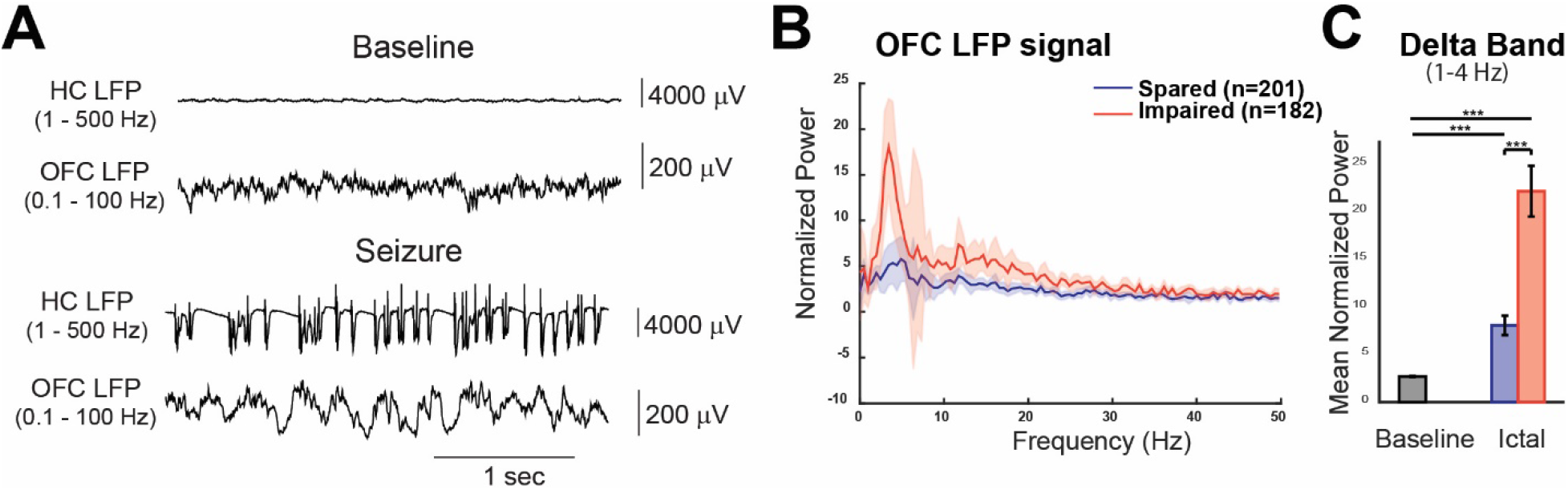
Increased cortical slow wave activity induced by focal limbic seizures is associated with impaired behavior. A. Example traces of slow wave activity recorded from orbitofrontal cortex (OFC) during induced seizure in hippocampus (HC), in comparison to fast rhythms recorded from OFC during awake behaving state (labeled as baseline). B&C. OFC LFP signals for ‘spared’ behavior (blue trace) or ‘impaired’ behavior (red trace), during ictal period. Power spectrum of ‘Impaired’ behavior showed larger amplitude in power for delta frequency (1-4Hz), which correspond to slow wave activity, compared to the power of ‘spared’ behavior (B). ‘Impaired’ delta power is significantly higher than ‘spared’ and ‘baseline', while ‘spared’ behavior is intermediary (C), showing that increased cortical slow waves induced by seizures correlate to behavioral impairment. Error bar shows +/-standard error mean. t test was use for 2 group comparisons, n=20 animals, ***: p<0.01.

## Conclusion

Here, we developed a novel awake mouse model of focal limbic seizures with similar key characteristics observed in human temporal lobe seizure: (i) seizures confined to the limbic region, (ii) slow wave activity resembling sleep in frontal cortex, (iii) impaired behavior associated with cortical slow waves. This new mouse model offers opportunities to investigate neuronal mechanisms related to behavior and arousal during seizures. To our knowledge, this is the first animal model demonstrating impaired behavioral responsiveness during focal limbic seizures, and relating impaired behavior to cortical neurophysiological changes (Sieu et al. 2020; Sieu et al. 2019; Sieu et al. 2018). Genetic tools available in mice (such as optogenetics or genetically encoded calcium indicators) offer more possibilities to explore neuronal networks and mechanisms (Cardin et al. 2010; Cardin 2012; Adamantidis et al. 2015; Crouse et al. 2020). Head-fixed mice provide good control of behavioral environment while allowing for simultaneous electrophysiologic recordings that would otherwise be difficult in a freely-moving animal. Overall, this mouse model of TLE opens new paths for investigating fundamental mechanisms responsible for loss of consciousness during focal limbic seizures.

